# Corticotropin-Releasing Factor in the Nucleus Accumbens Does Not Drive High Levels of Cocaine Consumption

**DOI:** 10.1101/2025.02.25.640140

**Authors:** L. Gordon-Fennell, R.D. Farero, J.D. Jones, L.S. Zweifel, P.E.M. Phillips

## Abstract

Uncovering the neurobiological processes underlying substance use disorder informs future therapeutic interventions. Prior research implicates the corticotropin releasing factor (CRF) system as a major player in a wide variety of substance use disorder-like phenotypes. However, the complexity of the CRF system in regard to brain region specific effects and experience-dependent changes in activity is poorly understood. Employing a cocaine self-administration paradigm that induces escalation of cocaine consumption in a subset of subjects, we investigated the role of CRF activity in the Nucleus Accumbens (NAc) in cocaine-taking patterns both before and after chronic cocaine experience. Our results showed that pharmacologically inhibiting CRF-R1 in the NAc did not reduce cocaine consumption following escalation and genetically deleting CRF-R1 from cells in the NAc did not prevent escalation. Overall, this suggests that any effect of CRF activity driving escalation or high levels of cocaine consumption is not through its actions on CRF-R1 in the NAc.

## INTRODUCTION

Substance Use Disorder (SUD) is a brain disease that is characterized by a plethora of behavioral and physiological phenotypes. The current epidemic of SUDs has a major toll on individuals and society today, with cocaine SUD specifically still having no current pharmaceutical interventions. Revealing the underlying neurobiology of SUD and its various phenotypes is a necessary step toward the development of effective therapeutics. The SUD phenotype characterized by increases in drug consumption over time – termed escalation – is an ideal candidate for employing harm-reductionist approaches to combat the toll of SUD. Preventing or reversing escalation would result in lower levels of drug consumption, decreasing the risk of overdose, loss of opportunity due to frequency of use, likelihood of infection, and more. We can model escalation of drug consumption using rodent self-administration under long-access (~6hr) conditions^1^.

Corticotropin-releasing factor (CRF) is a neuropeptide most known for its role in stress-related behaviors and has been widely implicated in a variety of SUD phenotypes^2^. Stressful stimuli, such as drug withdrawal, are believed to be a factor in escalation of drug consumption which may be a result of CRF activity^3,4^. However, CRF activity is not a monolith – its activity results in different behavioral and neurobiological effects based on brain region and prior experience of the subject^5–8^. Brain-region specificity of CRF activity in relation to cocaine escalation is poorly understood.

The nucleus accumbens (NAc) is a basal forebrain brain region that receives dense innervation from midbrain dopamine neurons. Dopamine activity, particularly in the NAc, is related to the rewarding and reinforcing properties of drugs of abuse^9–11^. Related to cocaine escalation, a decrement of the dopamine phasic response in the NAc to contingent delivery of drug-associated cues drives increases in cocaine consumption^12,13^. Systems that are activated during drug consumption and can modulate dopamine in the NAc would be key candidates for further understanding the neurobiology of SUD. The majority of drugs of abuse stimulate hypothalamic and extra-hypothalamic CRF release^3,14,15^. Increases in CRF-R1 expression has been shown to increase nicotine self-administration^16^. Additionally, previous literature demonstrates that CRF positively modulates dopamine activity in the NAc under certain conditions and this effect is modulated by cocaine experience and stress^5,7^.

In the current research, we dissect the brain-region and broad temporal specificity of the role of CRF-R1 in escalation of cocaine consumption. To investigate the role of NAc CRF-R1 in later stages of SUD-like phenotype, we utilized pharmacology approaches to block NAc CRF-R1 after undergoing an escalation paradigm and measured changes in cocaine consumption. To investigate the role of NAc CRF-R1 in the development of this SUD-like phenotype, we utilized a novel CRISPR-SaCas9 viral vector to knock-out CRF-R1s prior to cocaine self-administration. Overall, we demonstrate a lack of necessity of NAc CRF-R1 activity in the development and sustainment of cocaine escalation.

## METHODS

### Subjects

Female (200-250g) and male (250-350g) Wistar rats were procured from Charles River Laboratories and singly housed in a temperature and humidity-controlled environment with access to enrichment and *ad libitum* laboratory chow and water. Rats were allowed at least one week of acclimation to their new environment before undergoing any experimentation, such as surgery. The housing room was on a 12hr on/off light cycle (lights on at 7:00am), and all behavioral experiments were performed during the light cycle. All animal use and procedures were approved by the University of Washington Institutional Animal Care and Use Committee.

### Surgical Procedures

All surgical procedures were performed under isoflurane (1-5%) using aseptic technique. Alternating swabs of betadine and alcohol (3x/each) were used to sterilize surgical sites. Rats were given an NSAID (meloxicam, 1mg/mL/kg in 3mL of saline, s.c.) on the day of surgery and post-operative day 1. Rats were allowed at least one week of recovery in between surgeries, if undergoing multiple surgical procedures.

### Cannula Implantation

For cannula implantation, rats were affixed onto a stereotaxic frame (Kopf) and major landmarks were measured (lambda and bregma) and utilized to flatten the skull. Four or more skull screws were secured into the skull in areas surrounding the site of interest to help reinforce the headcap. Burr holes were drilled overlaying the site of interest (NAc, angle: 0°, AP: 1.3mm, ML: +/−1.3mm, DV: −7.2mm). We lowered the custom bilateral guide cannula to 1mm dorsal to the target (−6.2mm) and secured the implant to the skull with dental cement (guide cannula details; 26G, 2.5mm center-to-center distance, 7mm projection, purchased from Plastics One, cat: 8IC235G26XXC). To prevent contaminants, we inserted a dummy cannula with 0mm projection (Plastics One) into the guide cannula and secured a dust cap (Plastics One) onto the top of the guide cannula.

#### Viral Injection

For viral injection, rats were affixed onto a stereotaxic frame (Kopf) and major landmarks were measured (lambda and bregma) and utilized to flatten the skull. Burr holes were drilled overlaying the site of interest (NAc, angle: 0°, AP: 1.3mm, ML: +/−1.3mm, DV: −7.2mm). A Hamilton syringe (7105KH 5.0ul SYR (24/2.75”/3), ref#: 88000) containing the viral cocktail was lowered to 0.2mm ventral the site of interest (−7.4mm). After a 2-minute pre-injection wait period, we utilized a Micro4 Microsyringe Pump to inject the viral vector into the site of interest at a rate of 250nL/min for a total of 1µL (4 min injection). Over the course of the first minute of the injection, the Hamilton syringe was slowly raised to 0.2mm dorsal to the site of interest (−7.0mm) where it remained for the rest of the injection time. We waited 8 minutes post-injection before raising the Hamilton syringe out of the brain. Viral injections were a 1:8 viral cocktail of AAV1-Cre-GFP and either AAV1-CMV-SaCas9-sgCrhr1 (crhr1 KO) or AAV1-CMV-SaCas9-sgRNA (control). After both injections were completed, we sutured the scalp with 4-0 silk suture and applied betadine to the incision. For all behavioral experiments, there was a minimum of four weeks between viral injection surgery and the short-access baseline cocaine self-administration sessions. This time allowed for sufficient knock-out of the CRFR1s.

#### Catheter Implantation

Catheters were made in-house using: a custom 26G guide cannula with a 5mm up/bent projection on one side and a 6mm projection cut below the pedestal on the other side (Plastics One, cat: 8IC315GFLX03), Liveo laboratory tubing (cat: 508-001), and DAP silicone. Male catheters measured a total length of 10cm, and female catheters measured a total length of 8cm. Final catheter length, specifically the length inserted into the vein, was determined during surgery. Catheters were sterilized in betadine for at least 24hrs before implantation.

Catheter implantation surgery was always performed second (if multiple surgeries) to reduce attrition due to loss of patency. During surgery, two surgical sites were prepared: the rat’s back, the anterior and posterior to the scapulae, and the area surrounding the right peck. The catheter was placed with the access-port secured to the back in between the scapulae and the tubing going over the right shoulder and down into the jugular vein. We used 4-0 silk sutures to secure the catheter into onto the vein, secure the pedestal into place, and close incisions. We used a locking solution of 60% polyvinylpyrrolidone-40 solution in saline containing gentamicin (20mg/mL) and heparin (1,000units/mL) at the conclusion of the surgery and capped the port with an in-house fabricated dust-cap (PE20 tubing). The locking solution remained in the catheter for 7-9 days before daily flushing began (flushing solutions: 0.9% saline or heparinized saline (80units/mL)). Rats were allowed at least 7 days of recovery before beginning cocaine self-administration.

### Drugs and Viral Vectors

CP 154,526 (Sigma, cat: PZ0100) was prepared in aCSF with 1% DMSO and 5% Cremophor and injected bilaterally into the NAc (0.5µL/hemisphere) 20 minutes before a cocaine self-administration session. Antalarmin hydrochloride (Sigma, cat: A8727) was prepared in 0.5% (w/v) carboxymethylcellulose (Sigma, cat: C5678) saline solution and injected intraperitonially 80 minutes before a cocaine self-administration session^17^. Cocaine hydrochloride was dissolved into 0.9% sodium chloride (5mg/mL) and filtered into 10mL syringes. Cocaine hydrochloride was obtained through the NIDA Drug Supply Program.

The following viral constructs were manufactured by the University of Washington Center for Excellence in Neurobiology of Addiction, Pain, and Emotion’s Molecular Genetics Resource Core: AAV1-CMV-SaCas9-sgCrhr1, AAV1-CMV-SaCas9-sgRNA, and AAV1-Cre-GFP. Titer for all viruses range from 0.5 to 5×10^9^ particles.

### Cocaine Self-Administration

Cocaine self-administration (SA) began a minimum of 7 days post catheter implantation. Rats were handled for at least 1-3 days prior to starting behavioral training. Prior to each cocaine SA session, rats’ catheters were flushed with either saline or heparinized saline (80units/mL).

Cocaine SA was performed in modular operant chambers (Med Associates) equipped with two nose-poke ports (each with their own nose-poke light), a white noise generator, a tone generator, a houselight, and a liquid swivel mounted on a weighted balance arm. Syringe pumps were used to deliver cocaine at a rate of 5RPM. Upon the start of the session, the white noise and houselight would turn on, signaling availability of cocaine. One nose-poke port was designated the “active” port and the other nose-poke port the “inactive.” Active and inactive designations switched between cohorts to control for chamber side preferences, but a given designation was always the same for a specific rat. One operant response in the active port (FR1 schedule) resulted in an infusion of cocaine (0.5mg/kg/mL), the cessation of the discriminative stimuli for 20-seconds (white noise and houselight), and a 20-second audiovisual cue (nose-poke cue light and tone). This 20-second period was also the length of the time-out period during which additional operant responses into the active port were recorded but did not result in additional cocaine infusions. Operant responses in the inactive port were recorded but had no programmed consequence. Short-access sessions (ShA) were 1 hour in length and long-access sessions (LgA) were 6 hours in length.

Rats were trained on cocaine SA under daily ShA conditions. To reach criteria of learning the task, rats had to obtain 10 or more infusions per session for three consecutive sessions. Therefore, training was a minimum of three days. In the unlikely event that an animal could not reach this criterion or were not stable in their drug consumption above 10 infusions, they were dropped from the study. After training, rats began the ‘baseline ShA’ block, consisting of five days of ShA sessions (1x/day), followed by two five-day blocks of LgA (LgAb1 and LgAb2). When investigating reversal of escalation, rats continued LgA past LgAb2 for about five days (LgAb3). Between blocks, rats had 0-3 days off from cocaine SA. To classify rats as escalators or non-escalators, we performed a linear regression analysis on active responses in the first hour of a session across the sessions of baseline ShA, LgAb1, and LgAb2. Rats with a significant, positive regression were classified as escalators, and rats with a non-significant or significant, negative regression were classified as non-escalators, as previously described^12^.

### Microinjections

We microinjected CP-154,526 (2µg/µL) or vehicle bilaterally into the NAc (0.5µL/hemisphere) 20 minutes prior to a LgA cocaine SA session. Rats had one day of normal LgA cocaine SA in between injection days. For the microinjection, we equipped a kdScientific injector (model 210) with two Hamilton syringes (1801RN 10µL SYR (26s/2”/2), ref#: 84877). We used A bilateral injector with 1mm projection (Plastics One, part#: 8IC235ISPCXC) to insert into the guide cannula and waited for 2 minutes post-injection before removing the injector. The same setup was used to microinject Blue Sky dye during euthanasia to confirm injection placements.

### Histology

For rats that underwent neural manipulations (viral injections and intra-NAc microinjections), we euthanized with pentobarbital (150mg/kg) and performed transcardial perfusion of 0.9% sodium chloride followed by 4% paraformaldehyde. Their brains were removed, additionally post-fixed in 4% paraformaldehyde for 24 hours, and then submerged in ascending percentages of sucrose (ending with 30% sucrose in PBS). The brains were then frozen and sliced into 40µm slices on a cryostat. To determine the location of our neural manipulations, we analyzed the location of the blue dye and/or cannula implant for microinjections and the location of GFP fluorescence for viral manipulations.

### Immunohistochemistry

For the CRISPR *crhr1* KO validation, neural tissue was collected in the same manner as all other rats that underwent neural manipulations. After collection, brain sections containing the cannula implant track were used to perform immunohistochemistry against HA-tag (for the CRISPR virus) and c-fos. Fluorescence images were taken using an apotome microscope.

### Statistical Analysis and Data Visualization

Statistical analyses and data visualization was performed in R (The R Foundation). The following statistical analyses were performed with the corresponding package: Mixed effects model – “lme4”; ANOVA – “afex”; HSD – “emmeans”; linear regression – base R. In all plots, error bars represent standard error of the mean. Data visualization was performed with “ggplot2”. Data visualization and unpaired t-test for the CRISPR *crhr1* KO validation was performed in Prism. Adobe Illustrator (CS4) was used for final figure compilation.

## RESULTS

### Global CRF-R1 Activity Contributes to the Maintenance of High Levels of Cocaine Consumption

Previous work has demonstrated that systemic antagonism of CRF-R1 reduces cocaine consumption in male rats undergoing long access cocaine self-administration^17^. We validated this effect with our behavioral setup in male and female rats (Fig 1A). We trained rats on a baseline of 5 short-access cocaine SA sessions followed by 9-10 long-access sessions before being performing within-subjects systemic injection of a CRFR1 antagonist (antalarmin, 25mg/kg, s.c.) 80 minutes prior to a long-access session (Fig 1B). As a group, rats increased in their cocaine consumption across behavioral sessions as determined by a mixed effects model showing a main effect of session (Fig 1C). Systemic injection of antalarmin reduced the total cocaine consumption during the first hour of the task relative to vehicle injection day (Fig 1E). This effect appeared minor, with the change in mean active responses only a reduction 3.4 responses from vehicle to antalarmin injection days, which may not be biologically relevant change. Possibly, the small effect seen in the first hour could be masking a larger, more transient effect earlier in the task, such as would be the case if only the load-up phase was affected. To analyze this, we broke down the first hour active responses into ten-minute bins. As expected, rats consume the most cocaine in the first ten-minute bin (load-up) and then sharply reduce consumption to be maintained at lower levels (Fig 1F). While there is a main effect of injection drug, post-hoc analyses did not reveal any significant differences between antalarmin and vehicle days for any given ten-minute bin, similar to previously described^17^.

**Figure 1.**
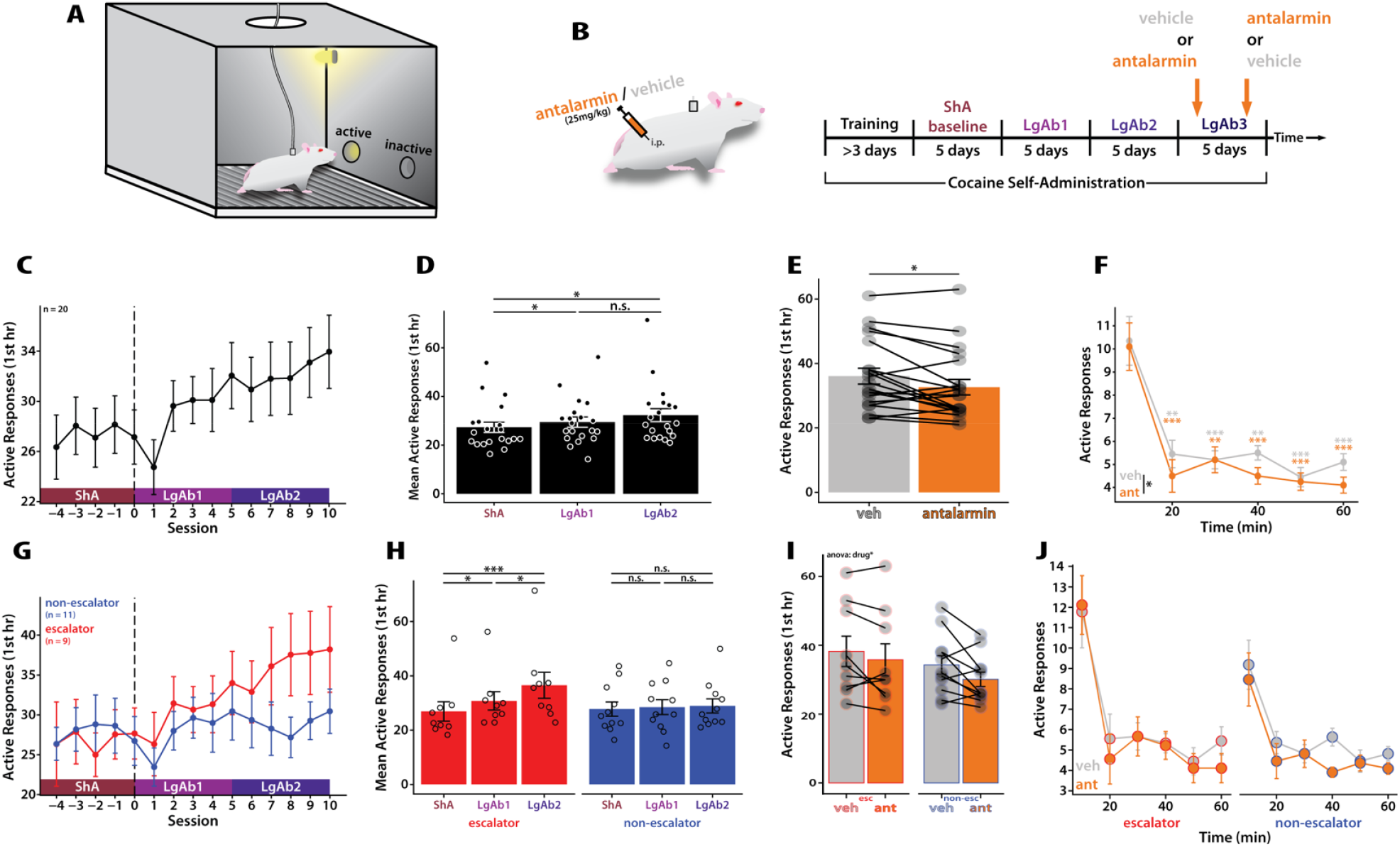
Systemic CRF-R1 Antagonism Reduces Cocaine Consumption Regardless of Escalation Status. **(A)** Behavioral setup. **(B)** experimental timeline. Antalarmin is a CRFR1 antagonist. **(C & G)** Number of operant responses in the active hole during the first hour of cocaine SA across behavioral sessions. **(C)** A mixed effects model showed drug consumption changes across behavioral sessions (main effect of session F(14,264)=5.23, p-value < 0.001). **(G)** A mixed effects model revealed a significant main effect of session (F(14,250) = 6.47, p-value < 0.001), no main effect of group (F(1,18) = 0.45, p-value = 0.51), and an significant interaction (F(14,250) = 3.24, p-value < 0.001). **(D & H)** Mean number of responses in active hole during the first hour of cocaine SA across behavioral blocks. (D) A repeated-measure ANOVA revealed a main effect of block (p-value < 0.001). **(H)** A repeated-measures ANOVA revealed a significant main effect of block (p-value < 0.001), a no main effect of group (p-value = 0.51), and a significant interaction (p-value < 0.001). **(E & I)** Number of operant responses in the active hole in the first hour of cocaine SA following vehicle and antalarmin injection.**(E)**A paired t-test revealed a significant difference between active responses in the first hour following vehicle vs antalarmin injection. **(I)** A repeated-measures ANOVA revealed a significant main effect of injection drug (p-value < 0.05) and no main effect of group (p-value = 0.21) nor interaction (p-value = 0.51). **(F & J)** Number of operant responses in the active hole in ten-minute bins across the first hour of cocaine SA for sessions following vehicle and antalarmin injection.**(F)** A repeated-measured ANOVA revealed a significant main effect of time (p-value < 0.001) and a main effect of injection drug (p-value < 0.05), but no interaction (p-value = 0.62). Statistics shown on plot are post-hoc Tukey of the given time bin compared to time bin = 10 for each injection drug. **(J)** A repeated-measures ANOVA revealed a significant effect of time (p-value < 0.001), injection drug (p-value < 0.05), and group X time (p-value < 0.05). All other comparisons were non-significant (group, p-value = 0.32; group X injection drug, p-value = 0.51; time X injection drug, p-value = 0.64; group X time X injection drug, p-value = 0.71). **(G-J)** Plots are grouped by ‘escalators’ vs ‘non-escalators,’ see methods for definition of escalation. **(all)** *** = p-value < 0.001; ** = p-value < 0.01; * = p-value < 0.05

The overall minor effect of antalarmin could also be a result of differential effects on different populations of rats. The ability of a CRF-R1 antagonist to reduce drug consumption in long-access paradigms is believed to be due to CRF activity underlying cocaine escalation. Therefore, we sought to determine if systemic CRF-R1 activity is necessary to sustain high levels of cocaine consumption selectively in rats that escalated in their cocaine consumption. Rats were classified as escalator or non-escalator based on a linear regression of their active responses in the first hour across behavioral sessions (see methods for more details). When grouping the rats by escalation status, we observed that escalators have a different pattern of cocaine consumption compared to non-escalators, where escalators overtake non-escalators by the end of LgAb2, as demonstrated by a significant interaction between session and escalation group (mixed effects model) (Fig 1G). Analyzing the behavioral data on injection days, while there is a main effect of injection drug, post-hoc analyses did not show a significant difference between antalarmin- and vehicle-treated sessions when rats were split up into their escalation group (Fig 1I). This lack of effect persisted when looking at ten-minute bins as well (Fig 1J). Similar lack of effects was observed when dividing by sex (Supp Fig 1). It is possible that the lack of effects seen when grouping the data by escalation group and/or sex is simply due to a larger number of animals being necessary to observe this seemingly biologically small effect.

### CRF-R1 Activity in the Nucleus Accumbens is Not Necessary to Sustain High Levels of Cocaine Consumption

To delineate a specific brain region underlying the effect seen with systemic CRF-R1 antagonism, we used the same behavioral paradigm and blocked CRF-R1 receptors only in the Nucleus Accumbens (NAc) through local microinjections of a CRF-R1 antagonist, CP 154,526 (Fig 2A). As a group, rats increased in their cocaine consumption across behavioral sessions as determined by a mixed effects model showing a main effect of session (Fig 2B). Local blockade of CRF-R1 in the NAc did not decrease cocaine consumption in the first hour (Fig 2D). Similarly, there was no significant differences between CP-and vehicle-treated days when the first hour of cocaine SA was broken into ten-minute bins (Fig 2E).

**Figure 2.**
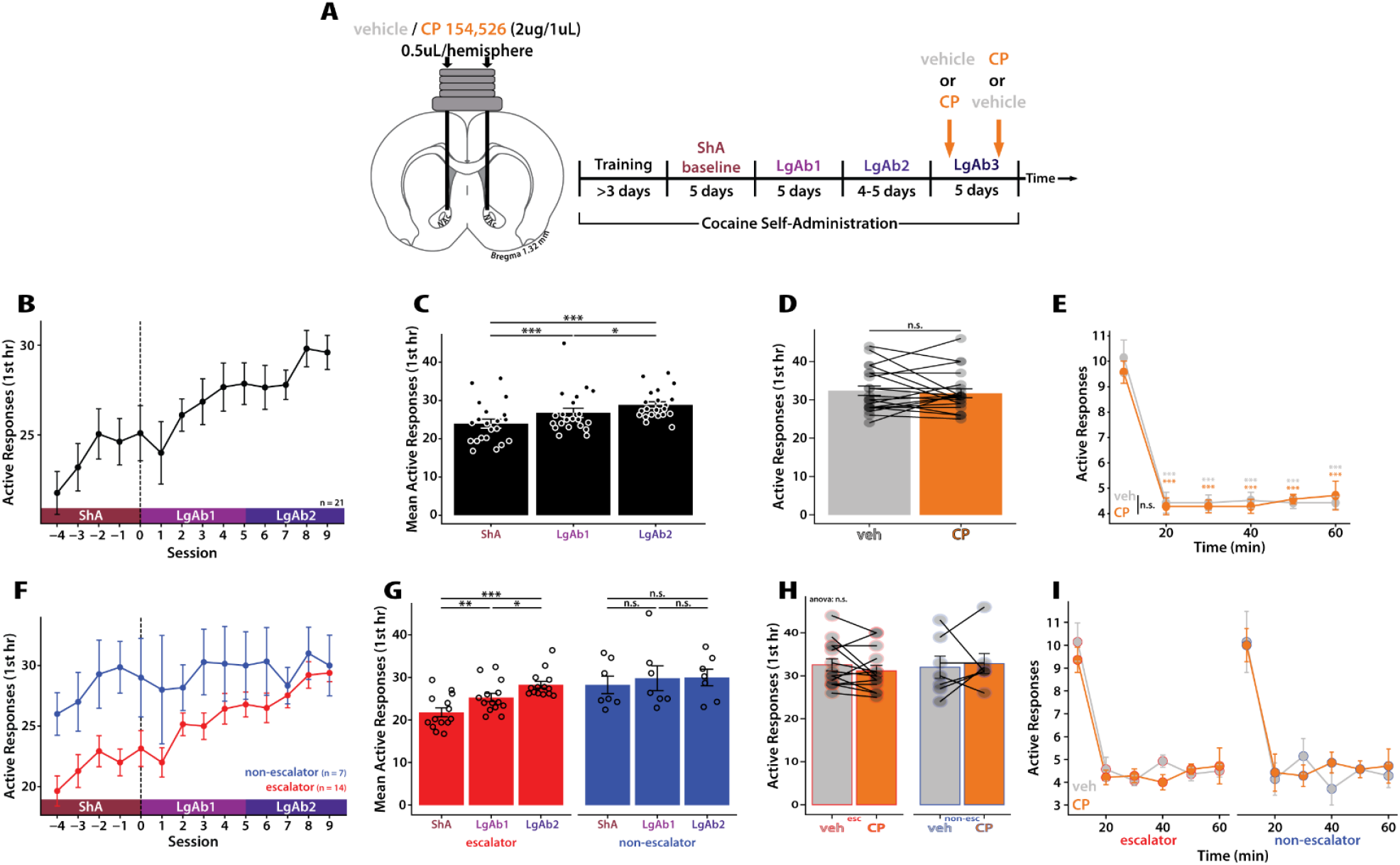
CRF-R1 Activity in the NAc is Not Necessary to Sustain High Levels of Cocaine SA. **(A)** Experimental timeline. CP 154,526 (CP) is a CRFR1 antagonist. **(B & F)** Number of operant responses in the active hole during the first hour of cocaine SA across behavioral sessions. **(B)** A mixed effects model showed drug consumption changes across behavioral sessions (main effect of session F(13,253)=9.89, p-value < 0.001). (F) A mixed effects model revealed a significant main effect of session (F(13,240) = 7.17, p-value < 0.001), main effect of group (F(1,19) = 4.74, p-value < 0.05), and an interaction (F(13,240) = 2.06, p-value < 0.05). **(C & G)** Mean number of responses in active hole during the first hour of cocaine SA across behavioral blocks. **(C)** A repeated-measure ANOVA revealed a main effect of block (p-value < 0.001). **(G)** A repeated-measures ANOVA revealed a main effect of block (p-value < 0.001), a main effect of group (p-value < 0.05), and an interaction (p-value < 0.01). **(D & H)** Number of operant responses in the active hole in the first hour of cocaine SA following vehicle and CP injection. (D) A paired t-test revealed no significant difference between active responses in the first hour following vehicle vs CP injection (p-value = 0.55). **(H)** A repeated-measures ANOVA revealed no significant effect of injection drug (p-value = 0.81), group (p-value = 0.81), nor interaction (p-value = 0.35). **(E & I)** Number of operant responses in the active hole in ten-minute bins across the first hour of cocaine SA for sessions following vehicle and CP injection. **(E)** A repeated-measured ANOVA revealed a significant main effect of time (p-value < 0.001), but no significant effect of injection drug (p-value = 0.56) nor interaction (0.82). Statistics shown on plot are post-hoc Tukey of the given time bin compared to time bin = 10 for each injection drug. **(I)** A repeated-measures ANOVA revealed a significant effect of time (p-value < 0.001) and all other comparisons non-significant (group, p-value = 0.81; injection drug, p-value = 0.81; group X injection drug, p-value = 0.35; time X injection drug, p-value = 0.86; group X time X injection drug, p-value = 0.31). **(F-I)** Plots are grouped by ‘escalators’ vs ‘non-escalators,’ see methods for definition of escalation. **(all)** *** = p-value < 0.001; ** = p-value < 0.01; * = p-value < 0.05

We sought to determine if there was a differential effect of intra-NAc CRF-R1 antagonism based on escalation status. Escalators and non-escalators showed different patterns of cocaine consumption across the sessions as demonstrated by a significant interaction between session and escalation group (mixed effects model). Non-escalators were stable in the cocaine consumption across sessions whereas escalators started low and increase their consumption over sessions (Fig 2F). When separated by escalation status, there was no effect of CP on cocaine consumption during the first hour of cocaine SA (Fig 2H) nor during any ten-minute bin (Fig 2I).

CRF-R1 Activity in the Nucleus Accumbens is Not Necessary for Escalation of Cocaine Consumption There is evidence to suggest that the CRF system may be altered by chronic cocaine exposure^7,8^ and therefore, even though NAc CRF-R1 activity was not necessary to sustain high levels of cocaine consumption following long-access, NAc CRF-R1 activity may still be necessary for the establishment of cocaine taking patterns. To reduce the activity of CRF-R1 for a sustained period, we developed a novel virally mediated CRISPR-SaCas9 targeting *crhr1* (the gene encoding for CRF-R1) to completely knock-out CRF-R1 in cells. To functionally validate this technique, we unilaterally knocked-out *crhr1* in the NAc of rats before microinjecting CRF, euthanizing the rats, and assessing neuronal activation via c-fos immunostaining (supp fig 2). The virally transfected hemisphere had a significantly smaller number of c-fos positive cells than the control hemisphere, demonstrated a reduction in the functionality of CRF-R1 in those cells.

To determine if CRF-R1 activation is necessary for the development of escalation of cocaine consumption, we knocked-out (KO) *crhr1* in the NAc prior to the start of cocaine SA (Fig 3A). As a group, there was no difference in cocaine consumption across behavioral sessions between control and *crhr1* KO rats as demonstrated by a lack of a main effect of group and interaction (mixed effects model) (Fig 3B). Performing linear regression analysis, both control and *crhr1* KO groups showed a significant, positive correlation – demonstrating escalation as a group (Fig 3D). Looking at individual rats and their escalation status, there was no change in the proportion of escalators versus non-escalators (Fig 3E). There were no sex differences in cocaine consumption (Supp Fig 3).

**Figure 3.**
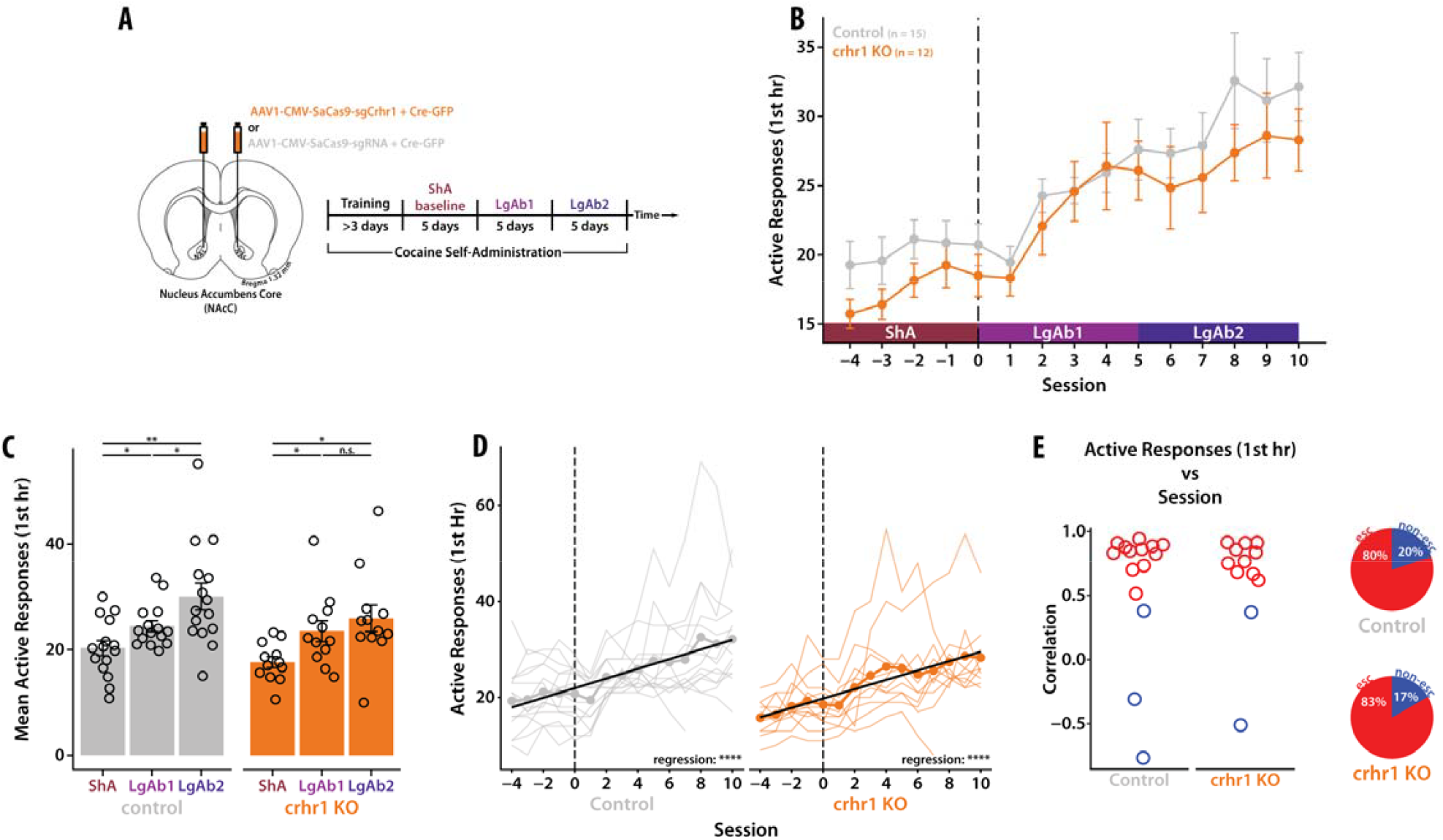
CRF-R1 Activity is Not Necessary for Escalation of Cocaine Consumption. **(A)** Experimental design. A minimum four-week incubation period occurred between viral surgery and ShA baseline. **(B)** Active responses in the first hour of cocaine SA across behavioral sessions. A mixed effects model revealed a significant main effect of session (F(14,339) = 19.02, p-value < 0.001), but not of group (F(1,25) = 1.17, p-value = 0.29) nor an interaction (F(14,339) = 0.46, p-value = 0.95). **(C)** Mean active responses in the first hour for each block. A repeated-measured ANOVA revealed a significant main effect of block (p-value < 0.001), but not of group (p-value = 0.25) nor an interaction (p-value = 0.49). (D) Same data as shown in B with the addition of individual subjects (thinner lines). The solid black line represents the best fit from the linear regression. Both control and *crhr1* KO groups showed a positive, significant regression demonstrating overall escalation of drug consumption as a group. **(E, left)** Correlation values (ρ) for individual subjects colored by escalation classification (red, escalator; blue, non-escalator). **(E, right)** Pie charts showing percentage of escalators vs non-escalators for control and *crhr1* KO groups. The proportion of escalators and non-escalators in the *crhr1* KO group is not significantly different from control (p-value = 0.82). **(all)** *** = p-value < 0.001; ** = p-value < 0.01; * = p-value < 0.05

## DISCUSSION

In alignment with previous literature^17,18^, we showed here that systemic antagonism of CRF-R1 reduces levels of cocaine consumption, albeit a very modest effect. However, this effect was not maintained when analyzing subgroups of rats based on escalation status or sex. Since CRF is known to have differential effects based on brain region, the small magnitude of this effect may be a result of simultaneously activating opposing systems via a systemic injection. Therefore, we sought to investigate specific brain region effects of CRF-R1 antagonism on cocaine consumption through local microinjection. We demonstrated that local CRF-R1 blockade in the NAc does not reduce cocaine consumption following long access cocaine SA. Considering that the effects of CRF, such as its ability to modulate dopamine in the NAc, are dependent on experience, we investigated the role of NAc CRF-R1 during an earlier timepoint of the task before the SUD-like phenotype of escalation emerges. We demonstrated that knock-out of CRF-R1 in the NAc does not reduce the magnitude of cocaine escalation nor the proportion of rats that escalate in their cocaine intake. Overall, this data suggests that CRF-R1 activation in the NAc is not necessary for the development or maintenance of escalation of cocaine intake.

CRF has long been implicated in SUD due to its involvement in for stress-induced changes in drug-related behaviors such as potentiation of drug reward^19^, reinstatement of drug seeking^20^, and potentiation of drug acquisition^21^. Additionally, CRF systems have been shown to be necessary for general negative affective behaviors resulting from stress and drugs of abuse^3,4,22–25^. These data position CRF to be a key player in the development or maintenance of the beta-process of opponent process theory. Elevations of CRF due to drugs of abuse or the stress of acute withdrawal could influence future drug taking behavior. Indeed, overexpression of CRF in the NAc leads to increases in CRF-R1 expression and nicotine SA in female rats^16^. However, the data presented here suggests that for cocaine, CRF-R1 activity in the NAc may not play a role drug taking patterns.

Appreciation for the diversity of the NAc and the roles of each subregion (such as: NAc core and NAc shell) has grown in the previous years^26^. The techniques employed here did not allow for subregion specificity. While our targeting mainly impacts the NAc core, the close proximity of the NAc shell and the changes in subregion medial-lateral and dorsal-ventral location throughout the anterior-posterior axis result in the considerable potential for volumes of the NAc shell to also be impacted. Therefore, similar to the systemic injection, these results may still be clouded by the manipulation of potentially opposing systems. Alternatively, the NAc shell has also been implicated in SUD and drug-taking behavior with relation to CRF activity^22,25,27^ and therefore, potentially targeting our manipulation more towards the NAc shell could reveal changes in cocaine consumption.

In this paper, we only investigated the role of CRF-R1. However, CRFR antagonists are known to not be entirely selective to CRF-R1 or CRF-R2 especially at high concentrations. Additionally, CRF-R2 have also been implicated in SUD and negative components of stress^4,16^. It is believed that the majority of CRF receptors in the NAc are CRF-R1, however, even with a smaller population of receptors, CRF-R2 could still play a role in modulating activity in the NAc to result in changes in drug-taking behavior^28^. Utilizing pharmacology with higher affinity to CRF-R2 and CRISPR targeting crhr2 would provide additional evidence of the role (or lack therefore) of NAc CRF system in cocaine consumption and SUD-related behaviors.

## ACKNOWLEDGEMENTS

We thank Dr. Olivier George for important discussion and advice, and to Hutch Clarke for assistance with histological analysis. This research was funded by the National Institutes of Health grants F31-DA059249 (LG), F31-DA048562 (RF), T32-DA007278 (PP & Ferguson), R25-DA057786 (Ferguson & PP) and R37-DA051686 (PP).

## SUPPLEMENTAL FIGURES

**Supp Figure 1.**
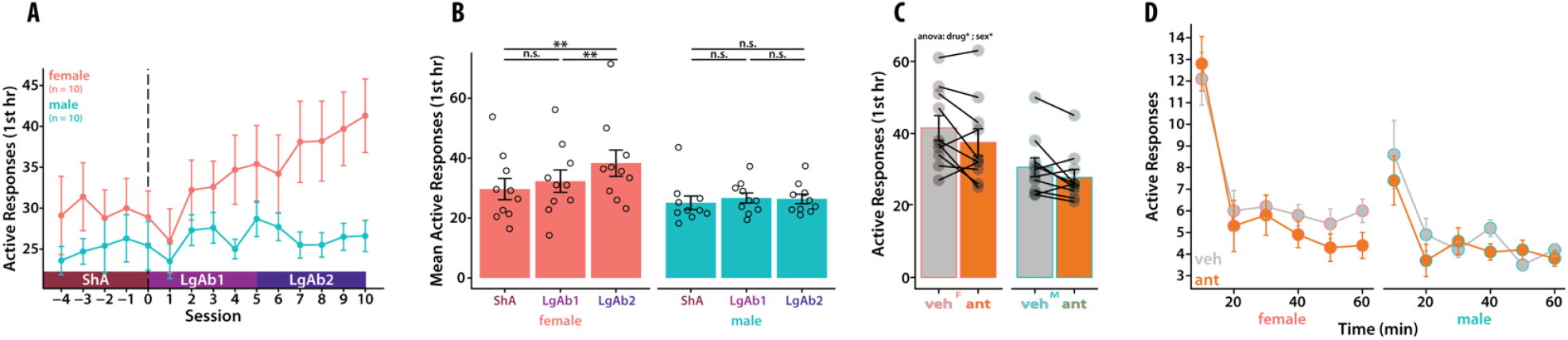
Antalarmin Effect on Cocaine SA in Males vs Females. **(A)** Number of active responses in the first hour of cocaine SA across sessions separated by sex. Mixed effects analysis revealed a significant main effect of session (p-value < 0.001) and a significant interaction (p-value < 0.001), but no main effect of sex (p-value = 0.09). **(B)** Mean number of active responses in the first hour of cocaine SA for each block separated by sex. A repeated-measured ANOVA revealed a significant main effect of block (p-value < 0.001) and a significant interaction (p-value < 0.01), but no main effect of sex (p-value = 0.09). **(C)** Number of active responses in the first hour of cocaine SA following vehicle and antalarmin (ant) injection. A repeated-measures ANOVA revealed a significant main effect of injection drug (p-value < 0.05) and sex (p-value < 0.05), but no interaction (p-value = 0.65). All relevant post-hoc Tukey were non-significant. (D) Number of operant responses in the active hole in ten-minute bins across the first hour of cocaine SA for sessions following vehicle and antalarmin injection separated by sex. A repeated-measures ANOVA revealed a significant main effect of time (p-value < 0.001), sex (p-value < 0.05), injection drug (p-value < 0.05), and interaction between sex and time (p-value < 0.01). All other comparisons were nonsignificant: sex X injection drug (p-value = 0.65), time X injection drug (p-value = 0.61), sex X time X injection drug (p-value = 0.20). **(all)** *** = p-value < 0.001; ** = p-value < 0.01; * = p-value < 0.05

**Supp Figure 2.**
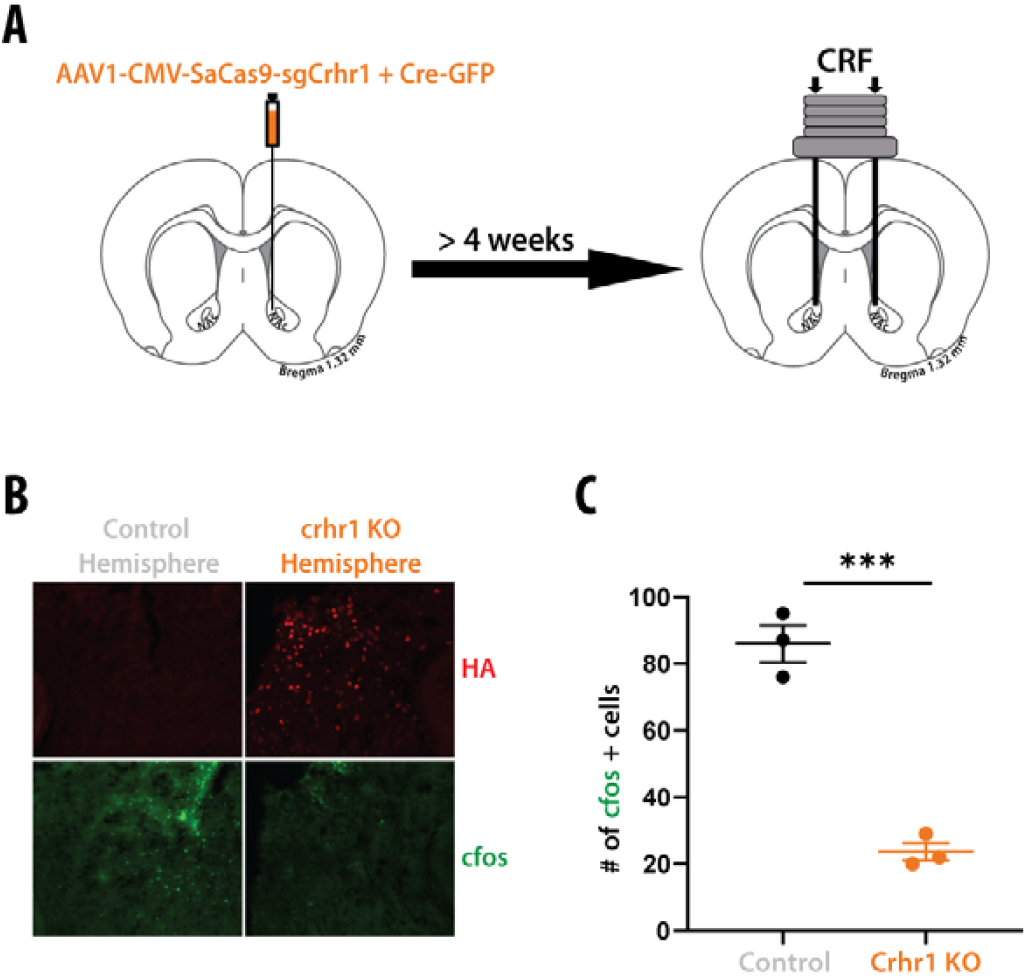
*crhr1* CRISPR Functionally Removes CRF-R1s. **(A)** Experimental Design. **(B)** Representative image of control and *crhr1* KO hemispheres immunostained against HA for the virus and cfos as a proxy for neural activity. **(C)** Number of cfos positive cells on control and *crhr1* KO hemisphere. Unpaired t-test (p-value < 0.001).

**Supp Figure 3.**
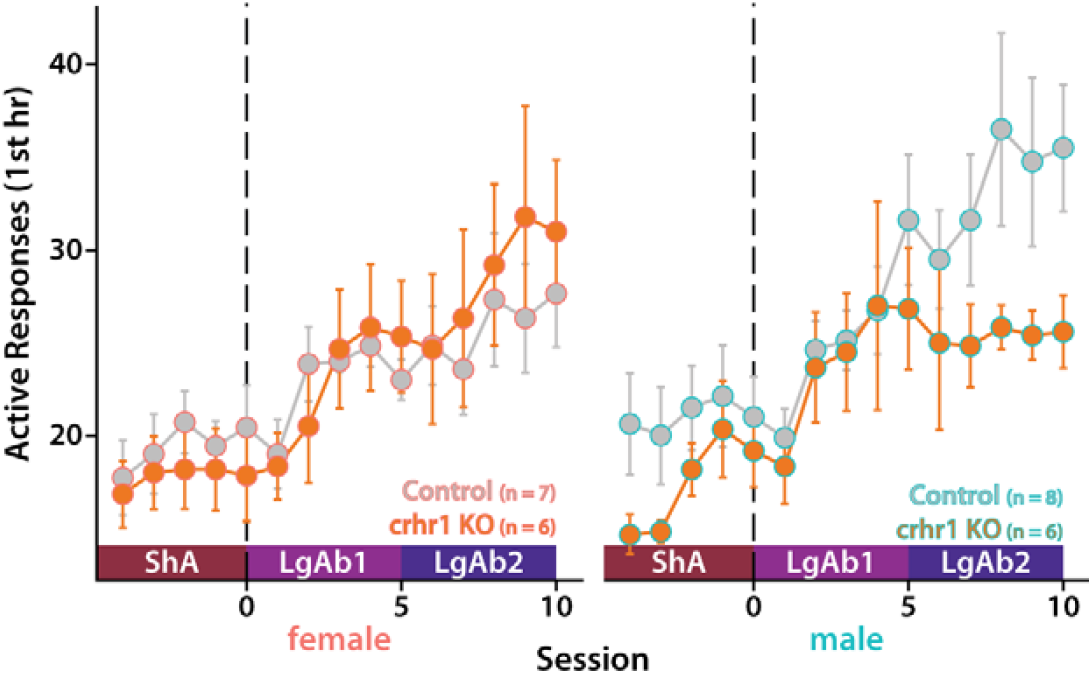
CRISPR KO of *crhr1* Effect on Cocaine SA in Males vs Females. Number of active responses in the first hour of cocaine SA across sessions separated by sex. Mixed effects analysis revealed no statistical significance for main effect of group or sex or any interactions.

